# Genetic Characterization of the *TAPBP* and Its Haplotypic Association with *BF2* in the Chicken Major Histocompatibility Complex

**DOI:** 10.64898/2026.04.20.719781

**Authors:** Roshani Fernando, Trisha Nicole Agulto, Eunjin Cho, Jaewon Kim, Andy van Hateren, Minjun Kim, Prabuddha Manjula, Jun Heon Lee

**Affiliations:** Department of Animal Science, Chungnam National University, Daejeon, 34134, Republic of Korea; Department of Animal Science, University of Peradeniya, Peradeniya, 20400, Sri Lanka; Centre for Immuno-Oncology, Nuffield Department of Medicine, University of Oxford, Oxford, OX3 7DQ, United Kingdom; Poultry Research Institute, National Institute of Animal Science, Rural Development Administration, Pyeongchang, 25342, Republic of Korea; Department of Animal Science, Uva Wellassa University, Badulla, 90000, Sri Lanka

**Keywords:** *TAPBP* (tapasin), MHC class I (*BF2*), Major Histocompatibility Complex (MHC), Co-evolution, Haplotype association, Avian Immunogenetics

## Abstract

*TAPBP* is a key chaperone of the peptide-loading complex that facilitates peptide loading onto major histocompatibility complex class I (MHC I) molecules. This study characterized *TAPBP* alleles in Korean Native Chickens (KNCs), identified novel variants, and evaluated haplotypic associations with *BF2*. Thirty-six samples representing six KNC lines were genotyped using LEI0258 and the MHC-B SNP panel, and individuals homozygous at both markers were classified into 16 groups. The same samples were subjected to Sanger sequencing of *TAPBP* exons 3-8. Sequences were assembled and aligned against MHC-B reference haplotypes and the Red Junglefowl reference. Additional comparisons with *“tapasin allele”* datasets enabled the identification of novel variants. Six novel nucleotide variants were detected across exons 3-6, including one nonsynonymous substitution in exon 4 (D251H). This residue corresponds to position Q265 in human *TAPBP* and lies adjacent to residues involved in MHC I interaction, suggesting potential functional relevance. Furthermore, *TAPBP* exhibited high haplotype diversity (Hd = 0.93) and moderate nucleotide diversity (π = 0.00892), with exon 5 showing the highest diversity (π = 0.01). B9 was the most frequent haplotype at the nucleotide level, whereas B6/B24 predominated at the amino acid level. Comparison with *BF2* data revealed haplotype-dependent pairing patterns: *BF2*-B9 consistently matched *TAPBP*-B9, whereas *BF2*-B6 was associated with distinct *TAPBP* nucleotide variants, indicating allelic diversification within a shared haplotypic background. Homozygosity at LEI0258 and the SNP panel corresponded with *TAPBP* homozygosity, supporting marker-based prediction. These findings highlight potential *BF2-TAPBP* associations and provide a foundation for understanding variation in MHC I peptide loading.

## INTRODUCTION

The chicken Major Histocompatibility Complex (MHC) is a compact, genetically diverse region encoding proteins central to adaptive and innate immunity, making it a critical focus for understanding avian immunity and genetic susceptibility to infectious diseases (Kaufman et al. 1999; Lamont, 1998; Silva & Gallardo 2020). The chicken MHC region consists of approximately 46 genes, spanning about 209 kb in the chicken genome (Silva & Gallardo 2020). Within the chicken MHC, there are class I, class II, class III, and extended class I regions (Kaufman et al. 1999; Lamont, 1998). The Class I region includes the extensively studied *BF1* and *BF2* genes, while the Class II region encompasses *BLB1* and *BLB2* genes. These Class I and II regions, known as the BF/BL region, contain 19 genes spanning a 92 kb region (Kaufman et al. 1999). Initial study of the B blood group of the chicken MHC (MHC-B) used serotyping, which revealed links between specific blood groups and their resistance or vulnerability to diseases. Variation in the MHC-B region is well-recognized for its role in conferring resistance to a range of highly pathogenic viral and bacterial diseases, as well as to internal and external parasites in poultry (Fulton et al. 2025; Jin et al. 2010; Kaufman et al. 1995; Lamont et al. 1987; Miller and Taylor 2016; Owen et al. 2008; Schou et al. 2010; Shiina et al. 2007). Efficient characterization of genetic diversity within the chicken MHC-B region requires reliable, time and cost-effective molecular markers. To address this need, initially, a high-density single-nucleotide polymorphism (SNP) panel was developed by Chazara et al. (2010) to identify the MHC-B diversity. Later, the panel was modified to 101 SNPs (currently used as the 90-SNP panel) by Fulton et al. (2016a, b) across different chicken lines as a time and cost-effective tool in identifying the genetic diversity of the MHC-B region along with the LEI0258 microsatellite marker (Esmailnejad et al. 2017; Manjula et al. 2020, 2021).

The chicken *TAPBP* (TAP-binding protein) gene, or tapasin, contains 8 exons and is located between the *BLB1* and *BLB2* genes within the class II region (Frangoulis et al. 1999; Kaufman et al. 1999). The *TAPBP* gene encodes a protein (tapasin) that plays a major role in the peptide loading complex (PLC) within the endoplasmic reticulum (Müller et al. 2022; Ortmann et al. 1997; Sadasivan et al. 1996). The PLC consists of the MHC class I (*BF1*/*BF2* and β2m), *TAP1* and *TAP2* heterodimers, calreticulin, and the ERp57 and the *TAPBP. TAPBP* assists MHC class I assembly in the endoplasmic reticulum by linking MHC class I molecules with the *TAP* heterodimer (Grandea et al. 1997; Li et al. 1997; Ortmann et al. 1997; Sadasivan et al. 1996; Solheim et al. 1997), stabilizing peptide-receptive MHC class I molecules at the site of peptide translocation, increasing the translocation of peptides via its association with *TAP* (Lehner et al. 1998), and editing the peptide repertoire (Howarth 2004; Wearsch and Cresswell 2007; Williams 2002) thereby enhancing the preferential loading of the high-affinity peptides onto MHC class I molecules. Among the 8 exons, exons 4 and 5 have been proposed to encode two luminal immunoglobulin (Ig)-like domains that retain characteristic cysteine residues and Ig-fold features, indicating they may play a key role as protein-protein interaction interfaces (Frangoulis et al. 1999; Williams and Barclay 1988). Furthermore, comparative sequence analyses indicate that exon 5 of the chicken *TAPBP* gene exhibits closer similarity to MHC class I and class II sequences, as reported by Frangoulis et al. (1999). However, subsequent structural studies of human *TAPBP* have refined this view. The first crystal structure revealed that the N-terminal region consists of a seven-stranded β-barrel fused to an Ig-like fold, forming a composite luminal domain that supports interactions within the PLC (Dong et al. 2009). Later structural and cryo-EM studies further demonstrated how *TAPBP* interacts with MHC class I and β_2_-microglobulin to facilitate peptide loading and editing (Jiang et al., 2022; Müller et al. 2022). Based on these structural insights, the domain encoded by exon 5, which lies proximal to the membrane and nestles between the α3 domain and β_2_-microglobulin, supporting the floor of the peptide binding domain in the crystal structures, is likely to play a critical role in MHC class I peptide loading. Based on mutagenesis studies conducted in mammals (Suh et al. 1999; Yu et al. 1999), two regions of the chicken MHC class I molecule have been proposed to interact with *TAPBP* (tapasin): the solvent-exposed loop in the α2 domain, formed by residues 125-133, and the solvent-exposed β strand in the α3 domain, formed by the residues 218-223. Furthermore, the importance of interactions between the membrane proximal domains of tapasin and MHC class I comes from studies of chicken MHC class I dynamics, where the α3 domain and β_2_-microglobulin exhibited altered conformational dynamics depending on peptide occupancy, highlighting their involvement in peptide loading and stabilization of the MHC class I complex (van Hateren et al. 2017).

Korean Native Chickens (KNC) are characterized by high genetic diversity and distinct molecular identity, slower growth and lower egg production compared with commercial broilers, and superior meat quality attributes such as firmer texture and enhanced flavor that drive strong consumer preference in South Korea (Choe et al. 2010; Haque et al. 2024; Jin et al., 2017; Jung et al. 2011). There are five KNC lines with distinctive feather colours: White (KNCW), Black (KNCB), Gray-brown (KNCG), Red-brown (KNCR), Yellow-brown (KNCY), and the heritage breed Yeonsan Ogye (YO). MHC-B diversity analysis (Manjula et al. 2020), NGS sequencing of the MHC-B region (Ediriweera et al. 2022), and identification of the MHC class I (*BF2*) haplotypes in the KNC populations provided an understanding of the disease-resistant and susceptibility haplotypes, and the findings could be incorporated into future breeding and conservation programs. Agulto et al. (2025) provided the first study to characterize the *BF2* diversity in KNC, having high polymorphism, in the peptide binding groove (α1 and α2 domains). These findings highlight the importance of MHC class I variation in shaping antigen presentation and immune responses. However, immune potential cannot be inferred solely from MHC class I polymorphism, as peptide loading also depends on *TAPBP*. Notably, *TAPBP* exhibits substantially greater polymorphism in birds, particularly in chickens compared with mammals (Halabi and Kaufman 2022; Kaufman 2015; Sironi et al. 2006), where only two alleles differing by a single amino acid have been reported in humans (Copeman 1998). van Hateren et al. (2013) reported that *TAPBP* and *BF2* co-evolve in a haplotype-specific manner, indicating coordinated evolution between these molecules to optimize peptide loading and MHC class I stability. Therefore, investigating *TAPBP* polymorphism is essential for understanding how variation in the peptide-loading machinery may contribute to disease resistance or susceptibility in KNCs.

Therefore, this study aimed to characterize the *TAPBP* alleles present in Korean Native Chickens (KNCs), identify novel sequence variants, assess whether individuals sharing identical *BF2* haplotypes also carry the corresponding *TAPBP* allele counterparts, and evaluate the association among LEI0258 allele sizes, BSNP haplotypes, and *TAPBP* alleles.

## MATERIALS AND METHODS

### Sample collection and DNA extraction

As described by Manjula et al. (2020), 379 samples were collected from 6 Korean native chicken (KNC) lines, including KNC Gray-brown (KNCG), KNC Black (KNCB), KNC Red-brown (KNCR), KNC White (KNCW), KNC Yellow-brown (KNCY), and the Yeonsan Ogye (YO). Genomic DNA was extracted from blood using the PrimePrep genomic DNA extraction kit (GenetBio, Daejeon, South Korea), followed by dilution for the downstream experiments: 5 ng/μL for the MHC-B SNP panel genotyping and 25 ng/μL for LEI0258 microsatellite marker genotyping and *TAPBP* sequencing.

### LEI0258 marker and MHC-B SNP panel genotyping

The samples were subjected to LEI0258 and MHC-B SNP panel genotyping to detect MHC-B diversity (Manjula et al. 2020). Samples that were homozygous for both the LEI0258 marker and MHC-B SNP panel were grouped into 16 groups (Agulto et al. 2025). In the present study, 36 homozygous samples were used for *TAPBP* gene sequencing: KNCG (*n* = 8), KNCB (*n* = 5), KNCR (*n* = 6), KNCW (*n* = 4), KNCY (*n* = 6), and YO (*n* = 7) (Table 1).

**Table 1.** Samples homozygous for both the MHC-B SNP panel and LEI0258 marker used for *TAPBP* gene sequencing.

| Group | BSNP Haplotype | Size of LEI0258 allele<br>(bp) | No. of Samples | Korean native chicken (KNC)<br>lines |
| --- | --- | --- | --- | --- |
| 1 | BSNP-Kr01 | 295 | 2 | KNCG |
| 2 | BSNP-Kr02 | 249 | 3 | KNCG |
| 3 | BSNP-Kr03 (rec) | 193 | 1 | KNCG |
| 4 | BSNP-Kr11 | 193 | 2 | KNCB |
| 5 | BSNP-Kr12 | 333 | 1 | KNCB |
| 6 | BSNP-Kr15 | 193 | 2 | KNCR |
| 7 | BSNP-Kr30 | 249 | 2 | YO |
| 8 | BSNP-Kr31 | 417 | 3 | YO |
| 9 | BSNP-J06 | 474 | 3 | KNCW |
| 10 | BSNP-T04 | 443 | 1 | KNCR |
| 11 | BSNP-R01 | 217 | 1 | KNCW |
| 12 | BSNP-O02 | 309 | 1 | KNCY |
| 13 | BSNP-Kr09 | 309 | 5 | KNCB (2)<br>KNCR (3) |
| 14 | BSNP-B03 | 249 | 3 | KNCG (1)<br>KNCY (2) |
| 15 | BSNP-T07 | 443 | 3 | KNCG (1)<br>KNCY (2) |
| 16 | BSNP-K02 | 249 | 3 | KNCY (1)<br>YO (2) |

*Note: KNC Gray-brown = KNCG; KNC Black = KNCB; KNC Red-brown = KNCR; KNC White = KNCW; KNC Yellow-brown = KNCY; Yeonsan Ogye = YO*

### *TAPBP* gene sequencing

Four sets of primers were designed spanning from exon 3 to 8 (only coding sequence) (Fig. 1). The PCR conditions were: 95 °C for 3 min, 30 -35 cycles of denaturation at 95 °C for 30 s, melting temperature (T_m_) as specified in Table 2 for 30 s, and extension at 72 °C for 15 to 45 s with a final extension at 72 °C for 10 minutes (Table 2; supplementary Table 1). Although several groups contained only a single individual, independent PCR amplification and sequencing were performed to confirm sequence accuracy and reproducibility.

**Table 2.**
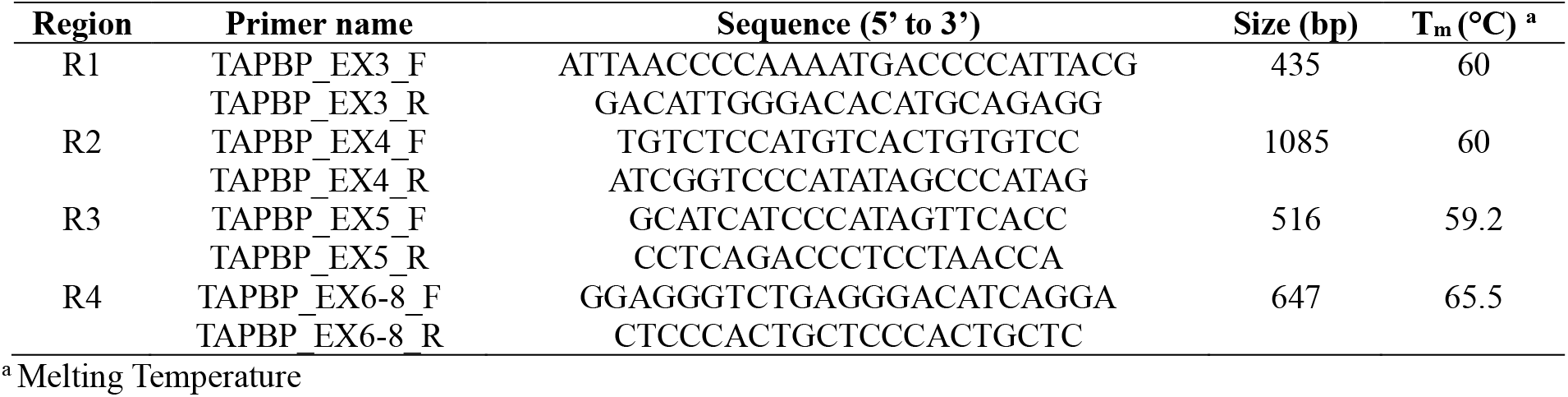
List of primers used to amplify the *TAPBP* gene.

**Fig. 1.**
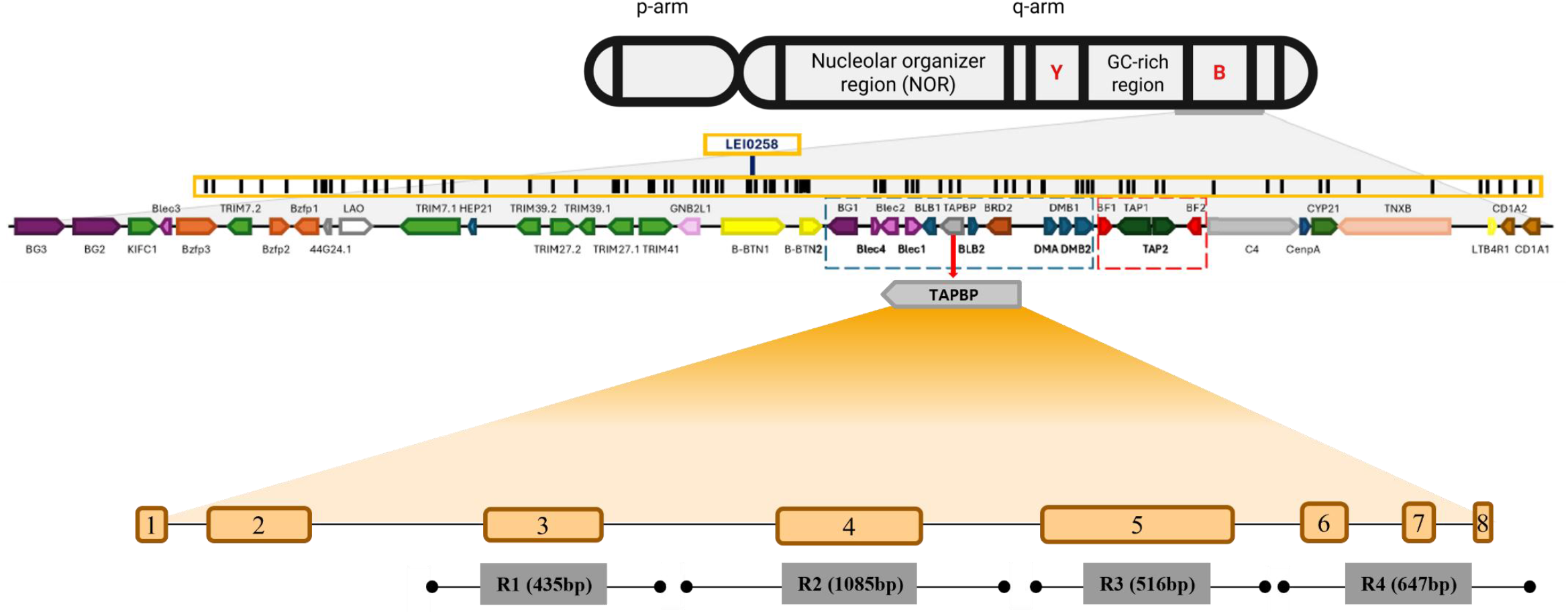
Genomic organization of the chicken MHC-B region on chromosome 16 and coverage of *TAPBP* gene sequencing Schematic representation of the MHC-B region on chromosome 16, indicating the relative positions of key genes, including *TAPBP*. The location of the LEI0258 microsatellite marker (Chazara et al. 2013) is highlighted (boxed), and black vertical lines denote the positions of the 90 SNPs comprising the MHC-B SNP panel described by Fulton et al. (2016a). Genes within the MHC-B region are indicated using different colored boxes. The enlarged lower panel illustrates the *TAPBP* gene structure, showing exons 1-8 and the regions targeted by four primer sets (R1-R4), which collectively span exons 3-8. Amplicon sizes for each primer set are indicated.

The PCR amplification products were confirmed by agarose gel electrophoresis and subsequently purified using the PrimePrep PCR Purification Kit (GenetBio, Daejeon, South Korea) before being subjected to Sanger sequencing (Macrogen, Sejong, South Korea). Sample sequences were submitted to GenBank (GenBank accession: PX685666 to PX685701).

## Data Analysis

### *TAPBP* diversity analysis

The obtained sequences were assembled using Geneious Prime software (v2024.0.7), and haplotype diversity (Hd) (using the ape, pegas, and seqinr packages) and the nucleotide diversity (π) (using the ape and pegas packages) were obtained through the R program v2024.12.0 (R Core Team, Vienna, Austria) before comparison with reference MHC-B haplotypes. This analysis provided an overall measure of genetic diversity within the study samples and supported downstream haplotype-based interpretation. Both indices range from 0 to 1, where values close to 0 indicate low diversity and values close to 1 indicate high diversity.

### Identification of *TAPBP* alleles and novel variants

The assembled *TAPBP* sequences were aligned against the 14 reference MHC-B haplotypes reported by Hosomichi et al. (2008), namely, B2, B5, B6, B8, B9, B11, B12, B13, B15, B17, B19, B21, B23, and B24, and a reference sequence from Red Jungle fowl (GenBank accession: AB268588) using Geneious Prime software (v2024.0.7) to identify *TAPBP* sequence matches to known haplotypes. Amplification and sequencing of exon 2 were unsuccessful, most likely due to the presence of extended homopolymer regions. Although exon 1 was successfully amplified and sequenced, sequence continuity across the gene could not be maintained; therefore, the subsequent analysis was focused on exons 3-8. Novel variants were identified through comparison with previously reported *“tapasin alleles”* (Tapasin*02, Tapasin*04, Tapasin*12, Tapasin*14, Tapasin*15, Tapasin*21) reported by van Hateren (2006), in addition to the reference sequences noted above.

### Association analysis between *BF2* and *TAPBP* alleles

Agulto et al. (2025) previously confirmed the presence of the standard B6 and B9 MHC-B haplotypes in the present sample set based on *BF2* gene genotyping. Building on this, the same individuals carrying B6 or B9 *BF2* haplotypes were used for a targeted assessment of whether they share specific *TAPBP* alleles, to explore potential haplotypic pairing patterns within the MHC-B region.

### Association analysis between LEI0258 marker allele sizes, BSNP haplotypes, and *TAPBP* alleles

The relationship between LEI0258 allele sizes, BSNP haplotypes, and *TAPBP* alleles was evaluated using a descriptive approach. Specifically, samples with identical LEI0258 allele sizes and BSNP haplotypes were evaluated to determine whether they also shared the same *TAPBP* allele.

## RESULTS AND DISCUSSION

### *TAPBP* diversity within the population

The haplotype diversity (Hd) of the *TAPBP* sequences was 0.93, indicating a high level of genetic variation among the samples studied. The overall nucleotide diversity (π) was 0.00892, and exon-specific analysis showed the highest nucleotide diversity in exon 5 (π = 0.010587), suggesting that sequence variation is relatively enriched in this region compared with other exons (Table 3). In contrast, exon 8 showed no detectable nucleotide diversity (π = 0), highlighting strong conservation. This exon-wise diversity pattern is consistent with previous reports (van Hateren, 2006), with exon 5 (339 bp) being the most polymorphic region and exon 8 (12 bp) relatively conserved. In humans, exon 5 encodes the membrane-proximal Ig-C domain of *TAPBP*, which is structurally and functionally implicated in interactions with MHC class I molecules and plays a key role in stabilizing peptide-receptive MHC class I complexes during peptide loading within the PLC (Dong et al. 2009; Müller et al. 2022). By contrast, exon 8 encodes part of the short cytoplasmic domain of *TAPBP*, for which no direct interaction with MHC class I or *TAP* has been clearly established. The marked difference in nucleotide diversity observed between exon 5 (most polymorphic) and exon 8 (highly conserved) likely reflects functional constraints across domains.

**Table 3.** Nucleotide diversity (π) and predicted protein region of each exon across exons 3-8 of the *TAPBP* gene in Korean Native Chickens (KNCs)

| Exon | Length of the exon (bp) | Nucleotide diversity ($\pi$) | Predicted protein region |
| --- | --- | --- | --- |
| 3 | 240 | 0.008472 | ER luminal domain |
| 4 | 345 | 0.009156 | ER luminal domain, Ig-V like domain |
| 5 | 339 | 0.010587 | Membrane proximal Ig-C domain |
| 6 | 90 | 0.004903 | Transmembrane domain |
| 7 | 26 | 0.006044 | Cytoplasmic domain |
| 8 | 12 | 0 | Cytoplasmic domain |

### Identification of *TAPBP* alleles in Korean Native Chickens (KNCs), based on reference MHC-B haplotypes

At the nucleotide level, the most frequent haplotype was B9 (22%), with eight samples exhibiting 100% identity to the MHC B9 reference haplotype (Table 4). This was followed by a grouped category comprising B5, B8, and B11 (17%; n = 6). At the protein level, the group category with highest similarity to B6 and B24 was the most prevalent haplotypes, followed by B9 (Table 4). Several samples showed complete correspondence to single reference haplotypes, including B9, B23, and B2 (Table 4). Notably, the B9 haplotype exhibited complete identity at both the nucleotide and protein levels. In contrast, Miller et al. (2004) reported 11 commonly occurring MHC-B haplotypes among 121 White Leghorn chickens, with B2, B5, B6, B7, B12, B13, B14, B15, B19, and B21 being most frequent, and several additional haplotypes occurring at lower frequencies. These differences are likely attributable to the limited sample size, the use of partial *TAPBP* exon 3-8 sequences, and differences in breed background. Moreover, because the MHC-B region comprises multiple tightly linked genes, the present analysis, restricted to a single locus (*TAPBP*), may reflect locus-specific variation rather than the overall distribution of complete MHC-B haplotypes. Supplementary Table 2 presents two identity matrices derived from the alignment of *TAPBP* nucleotide and amino acid sequences with reference haplotypes (Hosomichi et al 2008; van Hateren et al. 2013), showing the percentage similarity of the studied sequences to reference haplotypes other than the top-matching (highest similarity) haplotypes.

**Table 4.**
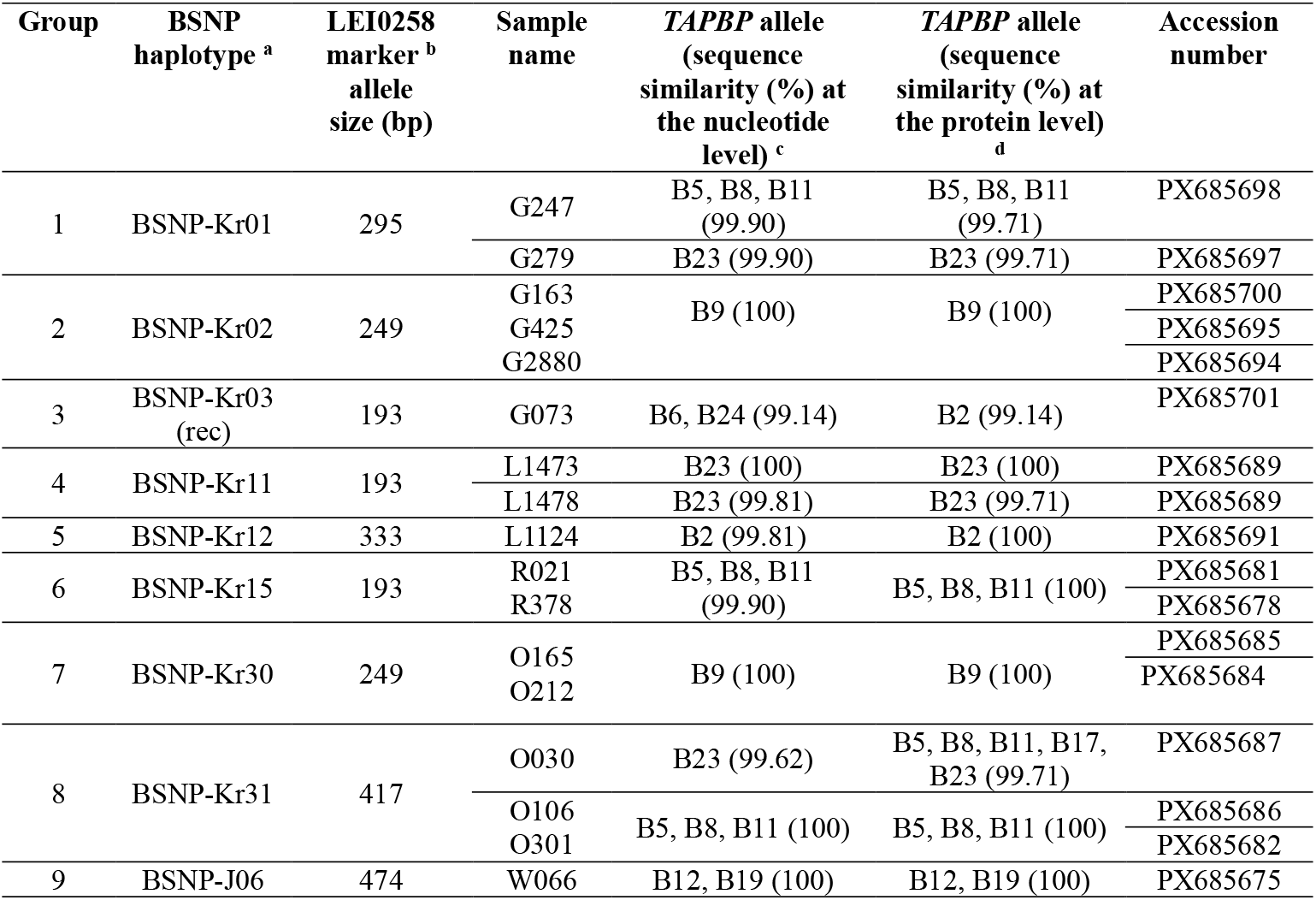
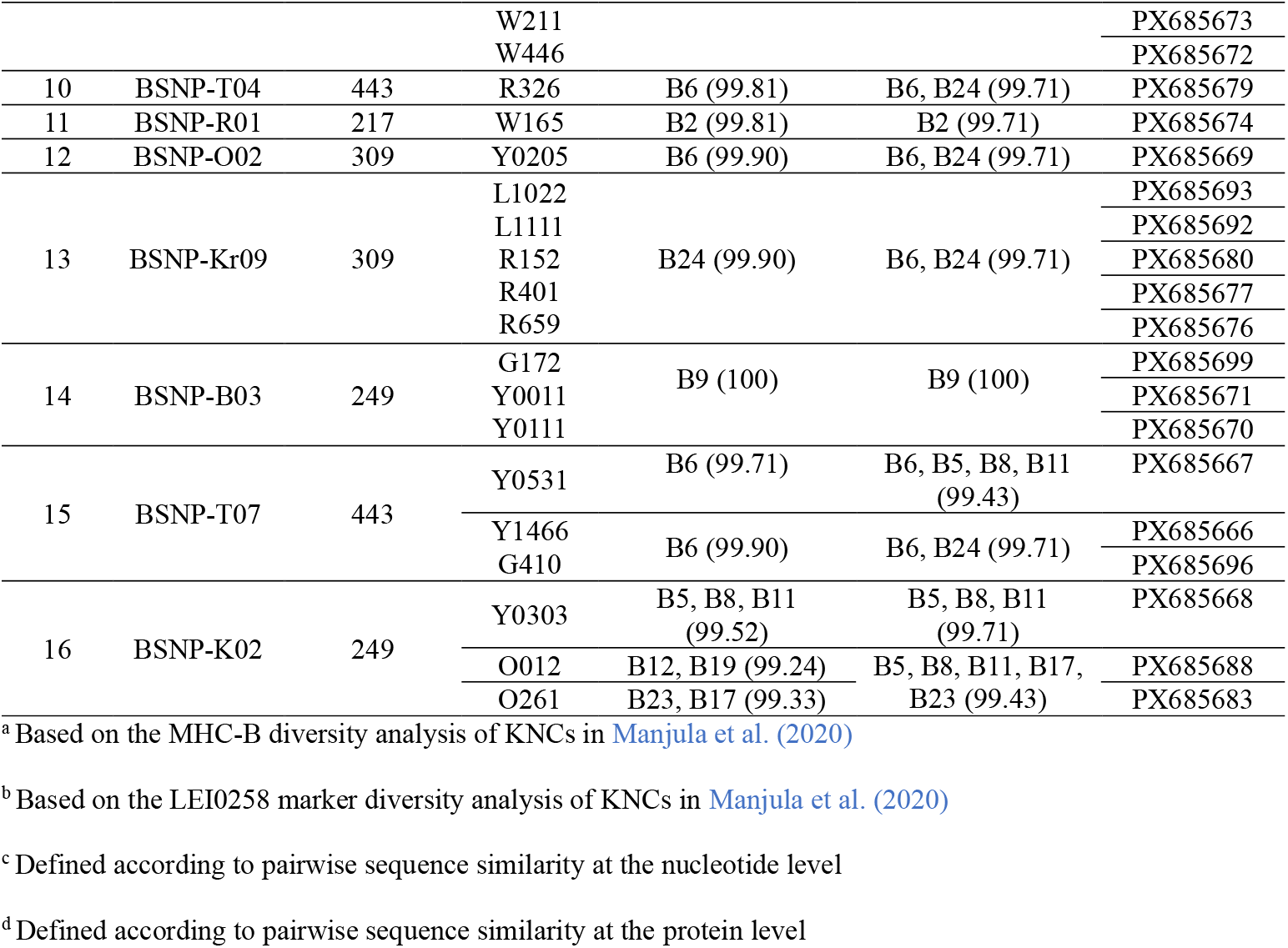
Categorization of *TAPBP* alleles at the nucleotide and amino acid levels based on reference MHC-B haplotypes defined by Hosomichi et al. (2008), using LEI0258 allele sizes and BSNP haplotypes. NCBI accession numbers for the deposited sequences are also provided.

### Nucleotide variants found in the *TAPBP* gene in KNCs

A total of 33 nucleotide variants were identified across the analyzed exons, with nearly three-quarters (n = 24) concentrated in exons 4 and 5 (Table 5a, b), consistent with previous findings (van Hateren, 2006). Of the total variants, six were novel, comprising five synonymous substitutions and one nonsynonymous substitution (Table 5a, b; Fig. 2). The highest number of novel variants was detected in exon 4. Exons 3 and 5 contained only a single novel synonymous variant and two synonymous variants, respectively, whereas no novel variants were observed in exon 7. Exon 4 included three novel variants, one of which was a nonsynonymous substitution (D251H) (Fig. 2). Notably, this novel nonsynonymous variant was observed in two samples in the KNC-Gray line. Comparative alignment of *TAPBP* sequences from multiple species (van Hateren, 2006) indicates that this position corresponds to Q265 in human *TAPBP* (Müller et al. 2022) (Fig. 3). Structural studies of human *TAPBP* have shown that residue Q262, located only three amino acids away, forms several hydrogen bonds with the α2-1 helix of MHC class I molecules, and mutation of this residue significantly reduced MHC class I surface expression (Müller et al. 2022). Although structural information for chicken *TAPBP* is currently limited, the proximity of the D251H substitution to a functionally important interaction region in the human ortholog suggests that this polymorphism may influence *TAPBP*-MHC class I interactions. Moreover, the substitution from Aspartate to Histidine represents a substantial change in physicochemical properties, which may alter interactions and potentially affect the stabilization of peptide-receptive MHC class I molecules.

**Table 5.**
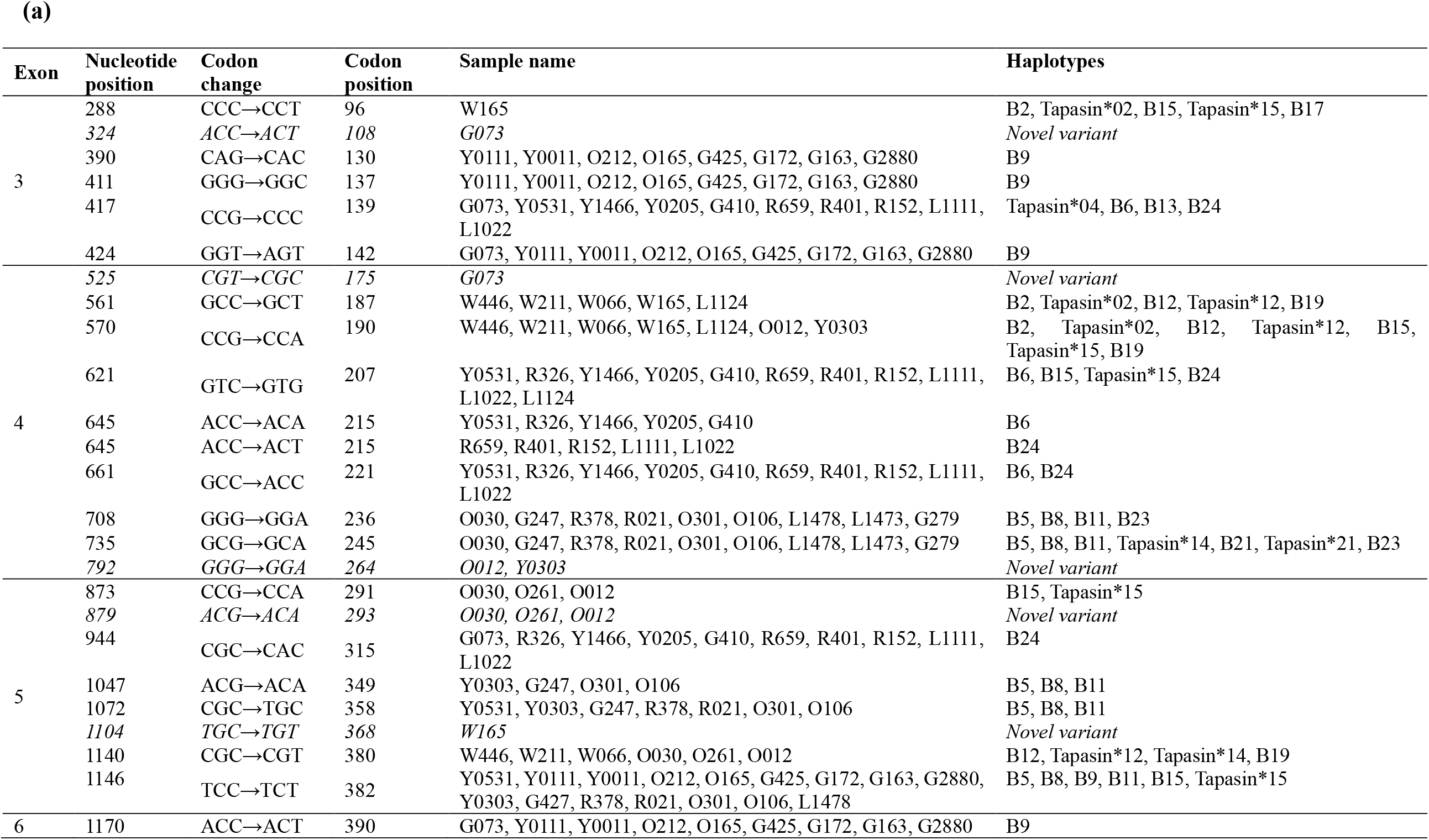
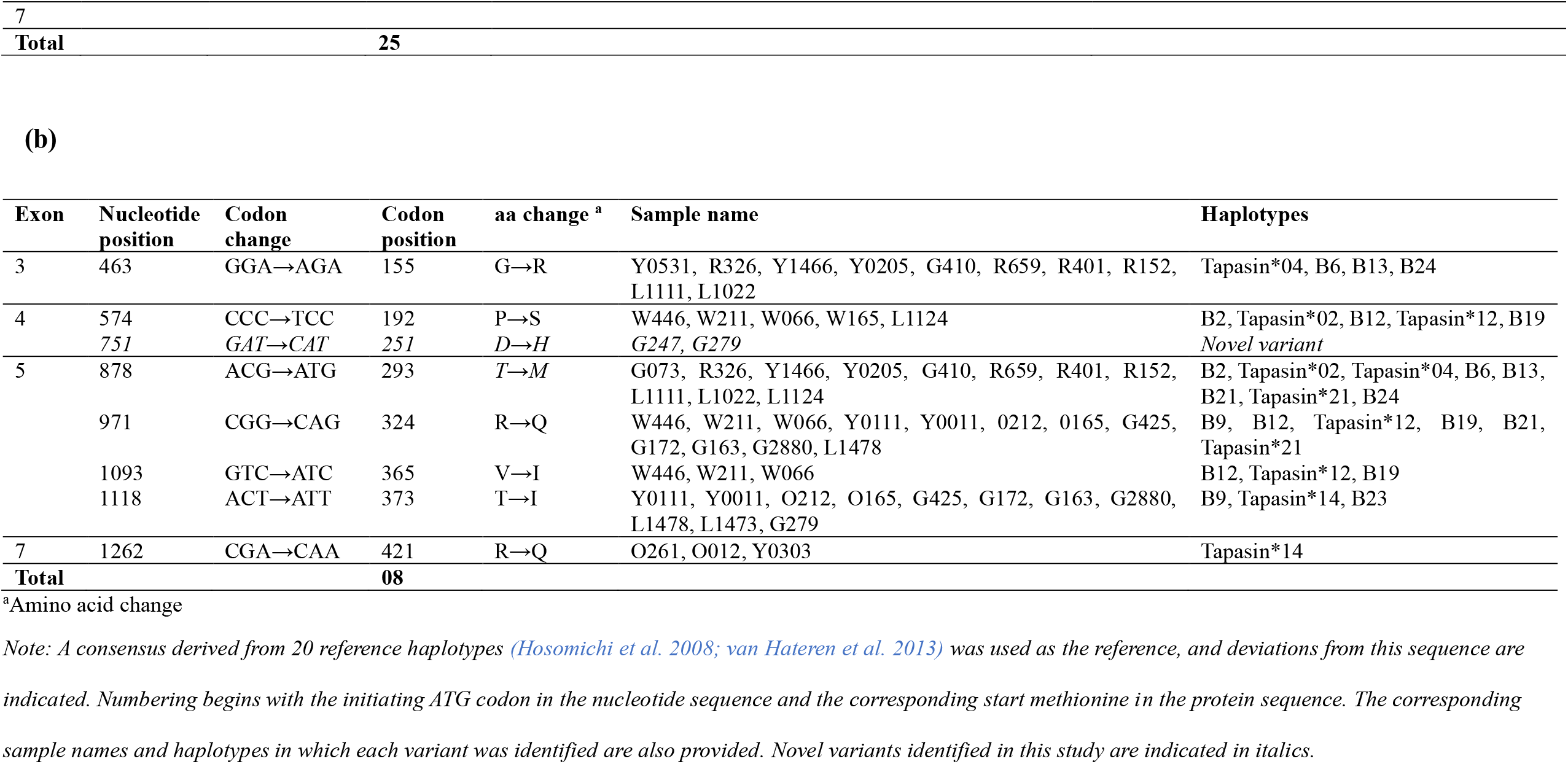
Nucleotide and amino acid polymorphisms identified in the *TAPBP* coding sequences relative to a consensus sequence. Variants are classified as (a) synonymous or (b) nonsynonymous substitutions.

**Fig. 2.**
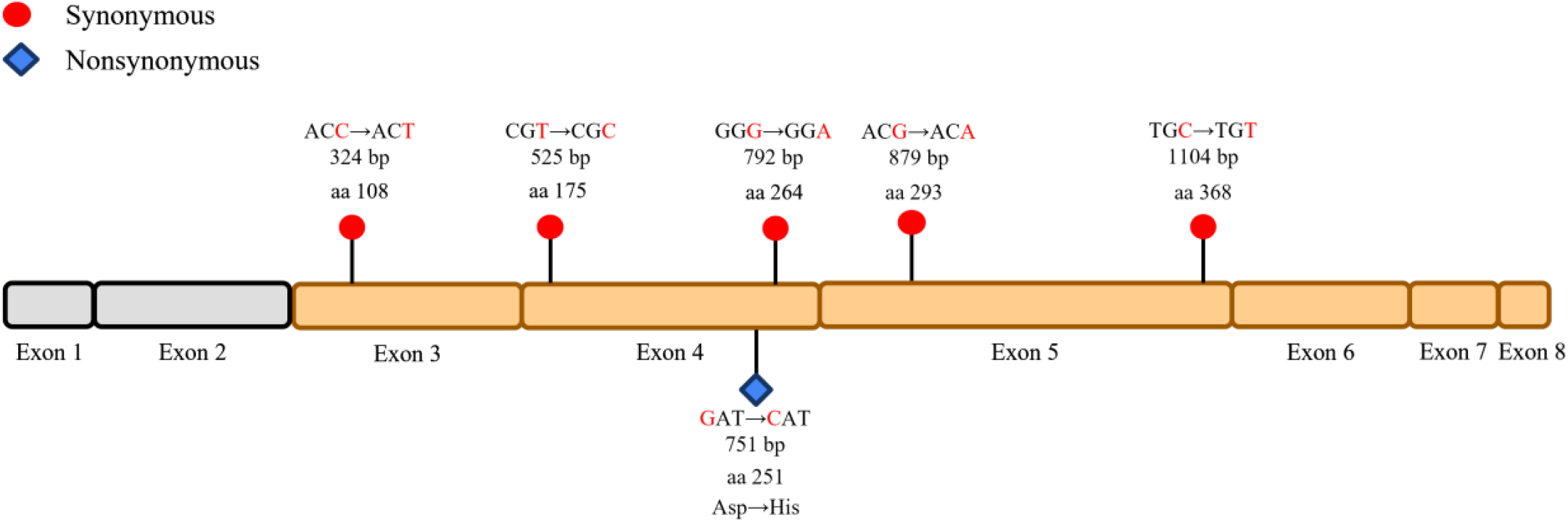
Distribution of novel nucleotide variants across exons 3-8 of the *TAPBP* gene in Korean native chickens (KNCs) Schematic representation of the *TAPBP* gene structure showing exons 1-8. Exons 1 and 2 are shaded in gray, indicating regions not included in sequencing. Novel nucleotide substitutions identified in exons 3-8 are indicated, with synonymous variants shown as red circles and the nonsynonymous variant as a blue diamond. Corresponding codon changes, nucleotide positions (bp), and amino acid substitutions (aa) are annotated above or below each Exon

**Fig. 3.**
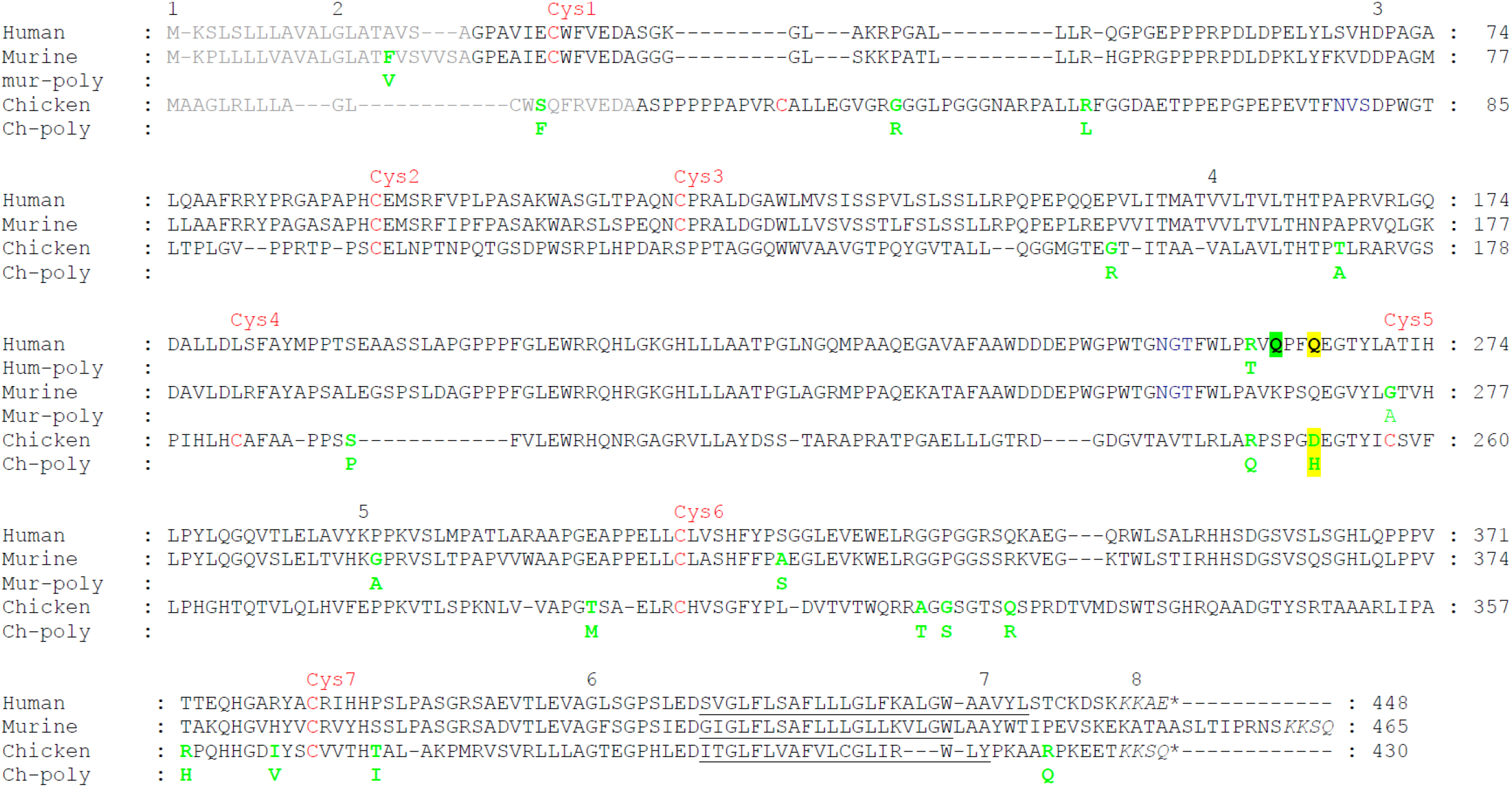
Alignments of human, murine and chicken tapasin protein sequences, with polymorphic amino acids indicated (adapted and modified from van Hateren, 2006). The alignment was generated in two steps: human and murine sequences were aligned following Grandea et al. (1997), and chicken sequences according to Frangoulis et al. (1999). Gaps were introduced to optimize alignment. Exon boundaries are indicated above the sequence. Dilysine motifs (italics), N-linked glycosylation sites (blue), predicted signal sequences (SignalP 3.0 program) (grey; human: Ortmann et al. 1997; murine: Grandea et al. 1997; chicken: this study; see also Frangoulis et al. 1999), and transmembrane domains are underlined (human: Ortmann et al. 1997; murine: Grandea et al. 1997; chicken: Frangoulis et al. 1999) are annotated. Conserved cysteine residues in ER luminal domains are shown in red. The novel chicken variant **D251H** identified in this study corresponds to **Q265** in human tapasin; the adjacent residue **Q262** is involved in MHC class I interaction (Müller et al. 2022). Polymorphisms identified in this study or reported previously are shown in green bold, with alternative residues indicated below (hum/mur/ch – poly). Previously reported polymorphisms include human position 260 (R/T) (Copeman et al. 1998), and murine tapasin positions 17 (F/V), 274 (G/A), 294 (G/A), and 326 (A/S) (Grandea et al. 1997; Li et al. 1999). Chicken polymorphisms identified across seven MHC haplotypes are indicated. Reference sequences used: human (AF009510), murine (AF043943; Grandea et al. 1997; differs from AF106278, Li et al. 1999), and chicken (AL023516; Kaufman et al. 1999)

[ufig]

### Association between *BF2* and *TAPBP* alleles

Chicken *TAPBP* is highly polymorphic and is thought to co-evolve with *BF2* within stable MHC-B haplotypes (Kaufman, 2020; van Hateren et al. 2013). Simonsen et al. (1980) reported that the linkage disequilibrium exists among MHC loci, however, it is not absolute over larger genomic spans, allowing differential allelic variation at distinct loci. Furthermore, Hosomichi et al. (2008) reported that recombination and gene conversion events within the MHC-B region can lead to allelic variation at linked genes even within conserved haplotypes. We found that in the B9 haplotype group, both *BF2* and *TAPBP* sequences showed 100% nucleotide identity with the corresponding B9 reference haplotype (Table 6). In contrast, within the B6 haplotype group, the *BF2* gene exhibited 100% nucleotide identity with the B6 reference haplotype, whereas the *TAPBP* gene displayed minor nucleotide differences (Table 6). This observation indicates that, despite identical classification at the MHC-B haplotype level based on *BF2*, minor nucleotide variations may exist within the *TAPBP* gene among individuals carrying the same B6 haplotype. Such variation suggests that the *TAPBP* gene may exhibit greater sequence flexibility relative to *BF2* within certain MHC-B haplotypes, potentially reflecting differences in selective constraints or recombination history within the region. However, to have a solid conclusion, the full *TAPBP* gene sequence is required for each haplotype, with a sufficiently large sample size.

**Table 6.**
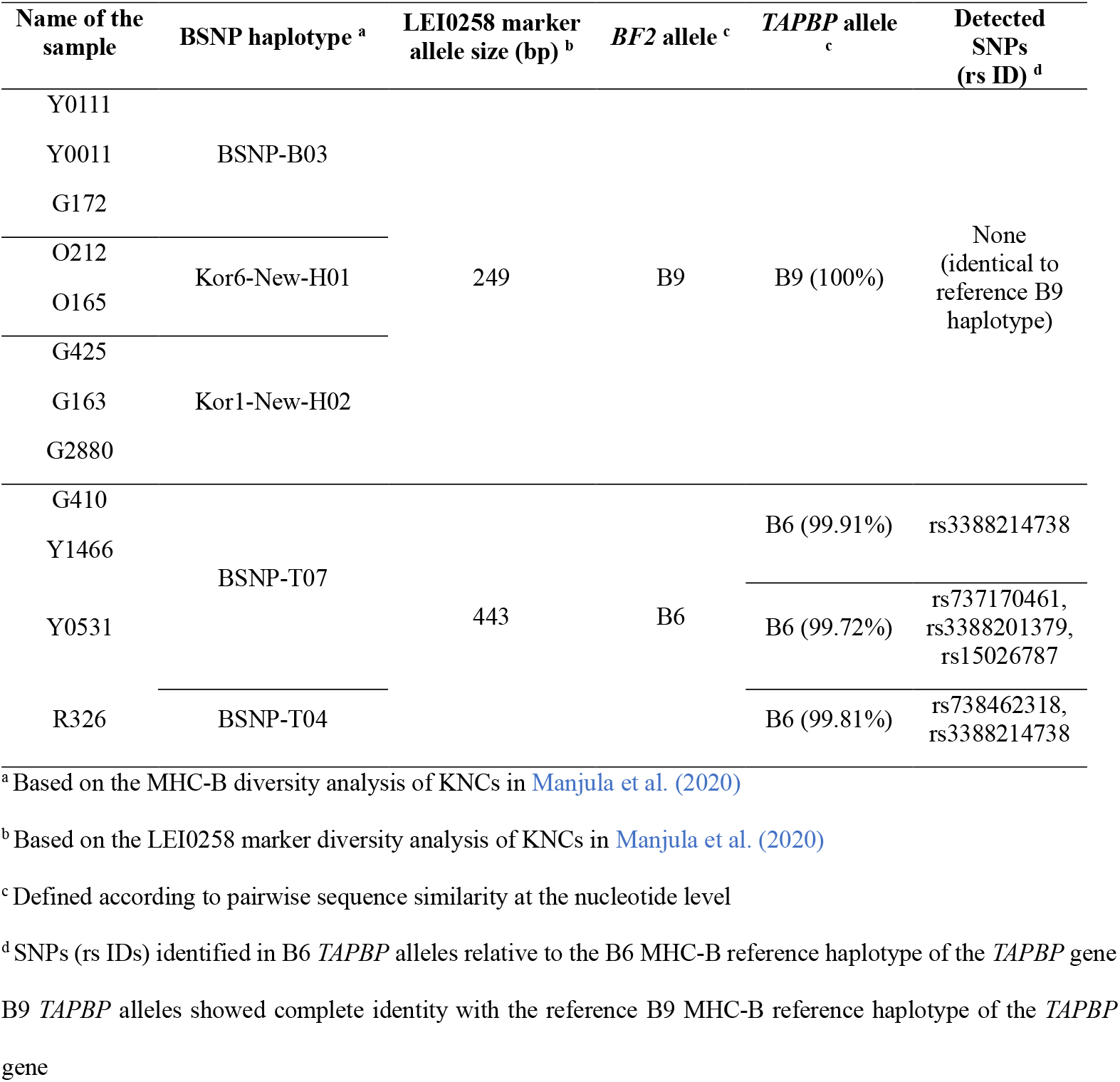
Association between *BF2* and *TAPBP* alleles observed within the B6 and B9 MHC-B haplotypes.

### Association among LEI0258 marker allele size, BSNP haplotypes and the *TAPBP* alleles

This study demonstrates that samples homozygous for both the BSNP haplotypes and the LEI0258 marker were also homozygous for the *TAPBP* gene sequences. These results are consistent with the findings of Agulto et al. (2025), who reported that samples homozygous for both the LEI0258 marker and BSNP haplotypes were also homozygous for the *BF2* gene in the MHC class I. Collectively, these findings validate the use of BSNP haplotypes and the LEI0258 marker as reliable indicators of homozygosity not only for *BF2* but also for the *TAPBP* gene.

In the present study, we identified 16 groups encompassing nine distinct LEI0258 allele sizes, ranging from 193 bp to 474 bp. At the CDS level, *TAPBP* alleles in groups 2, 4, 6, 7, 8, 9, 13, 14, and 15 were identical across all samples within each group (Table 4). At the protein level, identical *TAPBP* alleles were observed in groups 2, 4, 6, 7, 8, 9, 13, and 14 (Table 4). The LEI0258 249 bp allele size was consistently associated with the B9 haplotype, although one group exhibited a different MHC-B haplotype composition. Similarly, the B6 haplotype remained conserved across samples carrying the 443 bp LEI0258 allele size, despite variation in BSNP haplotypes. These observations are in agreement with Agulto et al. (2025), who reported that the 249 bp LEI0258 allele was predominantly associated with the B9 haplotype, with occasional sharing with B33, and that the B6 haplotype was consistently conserved among samples carrying the 443 bp allele. Consistent with earlier reports, Chazara et al. (2013) reported that the 443 bp LEI0258 allele is strongly associated with the B6 haplotype, while the 309 bp allele size is shared among multiple B haplotypes, including B10, B24, B26, and B76. Among these, the B24 haplotype was identified in the present dataset, except for one sample (Table 4).

### Limitations in sequencing the exon 2 region in the *TAPBP* gene

Several technical limitations were encountered during the sequencing of exon 2 of the chicken *TAPBP* gene, which consistently resulted in Sanger sequencing failure and apparent sequence variation. This was likely due to the exceptionally high GC content of exons 1 and 2 (73.8% and 77.4%, respectively) and the presence of extended homopolymer stretches in this region. Such sequence features are known to promote polymerase slippage during PCR and Sanger sequencing, consistent with the frame-shifted and overlapping chromatogram signals observed.

To evaluate whether the apparent variation represented true heterozygosity, cloning was performed; however, more than two alleles were obtained per individual, which is inconsistent with a diploid locus and suggests PCR-induced artefacts rather than genuine polymorphism. In addition, the same samples were previously genotyped using the LEI0258 marker and the MHC-B SNP panel (Manjula et al. 2020) and sequenced for *BF1* (Agulto et al. unpublished) and *BF2* (Agulto et al. 2025), all of which indicated homozygosity across the MHC-B region, making true heterozygosity in a short segment of *TAPBP* exon 2 unlikely.

Given the consistent technical difficulties associated with sequencing the GC-rich region spanning exons 1-2, these exons were excluded from the final analyses. For GenBank submission as a partial coding sequence (CDS), exon 1 was also omitted to maintain sequence continuity.

## CONCLUSION

In conclusion, the observed haplotype-dependent patterns suggest that *BF2-TAPBP* compatibility may vary among MHC-B haplotypes. Notably, the novel nonsynonymous substitution in exon 4 could alter *TAPBP* structure and/or interaction surfaces, potentially influencing its association with MHC class I molecules and, consequently, peptide loading efficiency and complex stability.

## ETHICS STATEMENT

The experimental method was approved by the Institutional Animal Care and Use Committee, the National Institute of Animal Science, Republic of Korea.

## CONSENT FOR PUBLICATION

All authors read and approved the manuscript.

## DATA AVAILABILITY STATEMENT

The sequences generated and analyzed in this study are available in the NCBI GenBank repository (GenBank accession: PX685666 to PX685701), with additional supporting data provided in supplementary files (Supplementary Table 1: Primer sequences and PCR profiles, Supplementary Table 2: Identity matrices of the *TAPBP* nucleotide and amino acid sequences).

## AUTHORS’ CONTRIBUTIONS

Conceptualization: Fernando R, Lee JH; Data Curation: Fernando R; van Hateren A; Formal Analysis: Fernando R; Methodology: Fernando R, Agulto TN; Software: Fernando R; Validation: Fernando R, van Hateren A, Manjula P, Lee JH; Investigation: Fernando R, Agulto TN, Cho E, Kim J, van Hateren A, Manjula P, Kim M, Lee JH; Writing (Original Draft): Fernando R; Writing (Review & Editing): Fernando R; Agulto TN, Cho E, Kim J, van Hateren A, Manjula P, Kim M, Lee JH.

